# Suppression of established invasive *Phragmites australis* leads to secondary invasion

**DOI:** 10.1101/2020.07.26.222018

**Authors:** C.D. Robichaud, R.C. Rooney

## Abstract

Invasive *Phragmites australis* (European Common Reed) is rapidly spreading throughout North American wetlands, with negative impacts on wildlife and native plants. The removal or suppression of *P. australis* is desired to provide an opportunity for native vegetation and wetland fauna to recover. In Ontario, managers applied a glyphosate-based herbicide to >400 ha of *P. australis* in ecologically significant Great Lakes coastal marshes, representing the first time this tool was used over standing water to suppress *P. australis* in Canada. Using a replicated Before-After-Control-Impact monitoring design, we 1) evaluated the efficacy of glyphosate-based herbicide at suppression *P. australis* along a water depth gradient and 2) assessed the recovery of the vegetation community for two years after treatment in relation to local reference conditions. We found that herbicide reduced live *P. australis* stem densities by over 99% the first year after treatment and about 95% the second year post-treatment. Treatment was equally effective along the entire water depth gradient (10 – 48 cm). The initial ‘suppression’ focused management was successful, but sustained monitoring and ‘containment’ focused follow-up treatment will be required to prevent reinvasion. Two years after treatment, the vegetation community does not resemble reference conditions. Although some treated plots initially increased in similarity to the vegetation communities typical of reference areas, many plots where *P. australis* was suppressed are on a novel trajectory comprising a vegetation community dominated by non-native *Hydrocharis morsus-ranae*. Secondary invasions represent a major challenge to effective recovery of native vegetation after *P. australis* control.

## Introduction

Globally, invasive species alter ecosystems in direct and indirect ways (Pyšek et al. 2020). The addition of one species can greatly modify community structure and ecosystem functions (Simberloff et al. 2013), and the majority of studied invasive plants negatively impact other plants at the species and community level (Pyšek et al. 2012). Theoretically, the removal of an invasive plant should catalyse the recovery of native species and ecosystem processes. Yet, well-established invasive species can become entrenched in the ecosystem, altering their environment (D’Antonio and Meyerson 2002), such that their removal can result in unanticipated ecological changes. In practice, although management often succeeds in suppressing the targeted invasive plant, secondary invaders are commonly the principal beneficiaries, and native species recovery is limited (Pearson et al. 2016). The removal of invasive plants can thus trigger undesirable outcomes that are difficult to anticipate (e.g., González et al. 2017). Unfortunately, invasive species treatment actions are rarely followed by adequate monitoring to provide accountability regarding invasive species management and native vegetation restoration outcomes (e.g., Blossey 1999; Kettenring and Adams 2011; Hazelton et al. 2014). For these practices to be considered successful, they should result in the recovery of native vegetation communities and not only the suppression of the invasive species that prompted the management action.

*Phragmites australis* ssp. *australis* ((Cav.) Trin. ex Steud.), a perennial grass introduced from Europe, is an aggressive invader of North American wetlands (Saltonstall 2002; Catling and Mitrow 2011). Once established, *P. australis* begins to change its environment. It produces extensive below and aboveground biomass (Lei et al. 2019), alters nutrient stocks (Yuckin and Rooney 2019) and creates a tall, dense canopy that shades out other wetland plants (e.g., Hirtreiter and Potts 2012), including species at risk (Polowyk and Rooney in prep). Unfortunately, because *P. australis* is so widespread in North American wetlands (Catling and Mitrow 2011; Carson et al. 2018), management of established populations is restricted to on-going asset-based protection and containment. For example, a survey of 285 U.S. land managers by Martin and Blossey (2013) found managers spent >$4.6 million USD/y on *P. australis* suppression. In Ontario, municipalities reported spending $2.8 million CAD to manage *P. australis* in 2019 alone (Vyn 2019). Given the large amount of public funds directed to *P. australis* suppression, it is critical that the efficacy of management actions is evaluated.

The application of either glyphosate- or imazapyr-containing herbicide is the most common management action applied to *P. australis* in North America (e.g., Martin and Blossey 2013; Hazelton et al. 2014; Hunt et al. 2017). Most studies that used herbicide report success in reducing the abundance of *P. australis* locally, but none suggest eradication was achieved. Some report short-term reductions in *P. australis*, and most caution that repeated control measures are needed to suppress re-growth (e.g., Lombard et al. 2012; Quirion et al. 2017). Fewer suggest that herbicide application led to the recovery of native vegetation. For example, Martin and Blossey (2013) report that although 75% of respondents agreed that management provided temporary suppression of *P. australis*, only 50% perceived an increase in the abundance of native plant species. To assess the response of native vegetation communities to different control methods, Rohal et al. (2019) contrasted the efficacy of *P. australis* management approaches, monitoring six treatment combinations over four years post-treatment. None of their treatments converged on the reference condition and native vegetation recovery was hydrology dependent. Greater recruitment of native species was reported in sites with higher soil moisture content, thus indicating the important role that hydrology plays in vegetation community assembly. Unfortunately, few studies evaluating *P. australis* management actions report on vegetation recovery in much detail (Hazelton et al. 2014).

We contend that *P. australis* treatment efficacy requires two elements: first, the treatment must reduce the stem density of *P. australis*; second, the treatment must lead to the recovery of native vegetation. Ideally, recovery should target a vegetation community that resembles the community present in equivalent edaphic and hydrologic conditions where *P. australis* never invaded (i.e., the reference condition, *sensu* Stoddard et al. 2006). Management outcomes for *P. australis* suppression techniques have received extensive study (e.g., Farnsworth and Meyerson 1999; Ailstock et al. 2001; Derr 2008; Mozdzer et al. 2008; Lombard et al. 2012; Rapp et al. 2012; Hunt et al. 2017; Quirion et al. 2017; Zimmerman et al. 2018; Judd and Francoeur 2019; Rohal et al. 2019a, b). Yet there are inconsistencies in the literature regarding information on baseline conditions, experimental controls, or adequately characterized recovery targets, making differences in outcomes difficult to reconcile.

The purpose of our study was to assess the efficacy of *P. australis* control in biodiversity hotspots on the north shore of Lake Erie, including two provincial parks, one of which is designated a World Biosphere Reserve, a Ramsar wetland, and an Important Bird Area. First, we determined how effective the aerial application of a glyphosate-based herbicide was at suppressing *P. australis* when applied over standing water of varying depths (10 – 48 cm). Second, we assessed the initial recovery of vegetation in the first two years post-treatment along the same water depth gradient, comparing recovering vegetation to uninvaded, reference emergent and meadow marsh vegetation communities. We used a spatially replicated BACI design with control and treatment plots paired by water depth to allow us to quantify the influence of water depth on treatment efficacy.

## Methods

### Study Area

Our study took place in two marsh complexes located on the north shore of Lake Erie: the Long Point peninsula and Rondeau Provincial Park. These Great Lakes coastal marsh complexes are approximately 165 km apart and are directly connected to their respective bays and sheltered from Lake Erie proper by sand bars (Fig. 1). Long Point and Rondeau represent over 70% of the remaining intact wetlands on the north shore of Lake Erie, and as such provide habitat for a number of rare and at-risk species (Ball et al. 2003). This ecosystem is threatened by *P. australis. Phragmites australis* abundance is well documented in Long Point, where it spread exponentially in the late 1990s, likely due to the introduction of the invasive haplotype and low water levels (Wilcox et al. 2003). Of the 217 species designated as Species at Risk under Ontario’s Endangered Species Act (S.O. 2007 c.6), 25% are directly threatened by invasive *P. australis*, including 24 species of vascular plants (Bickerton 2015).

**Figure 1.**
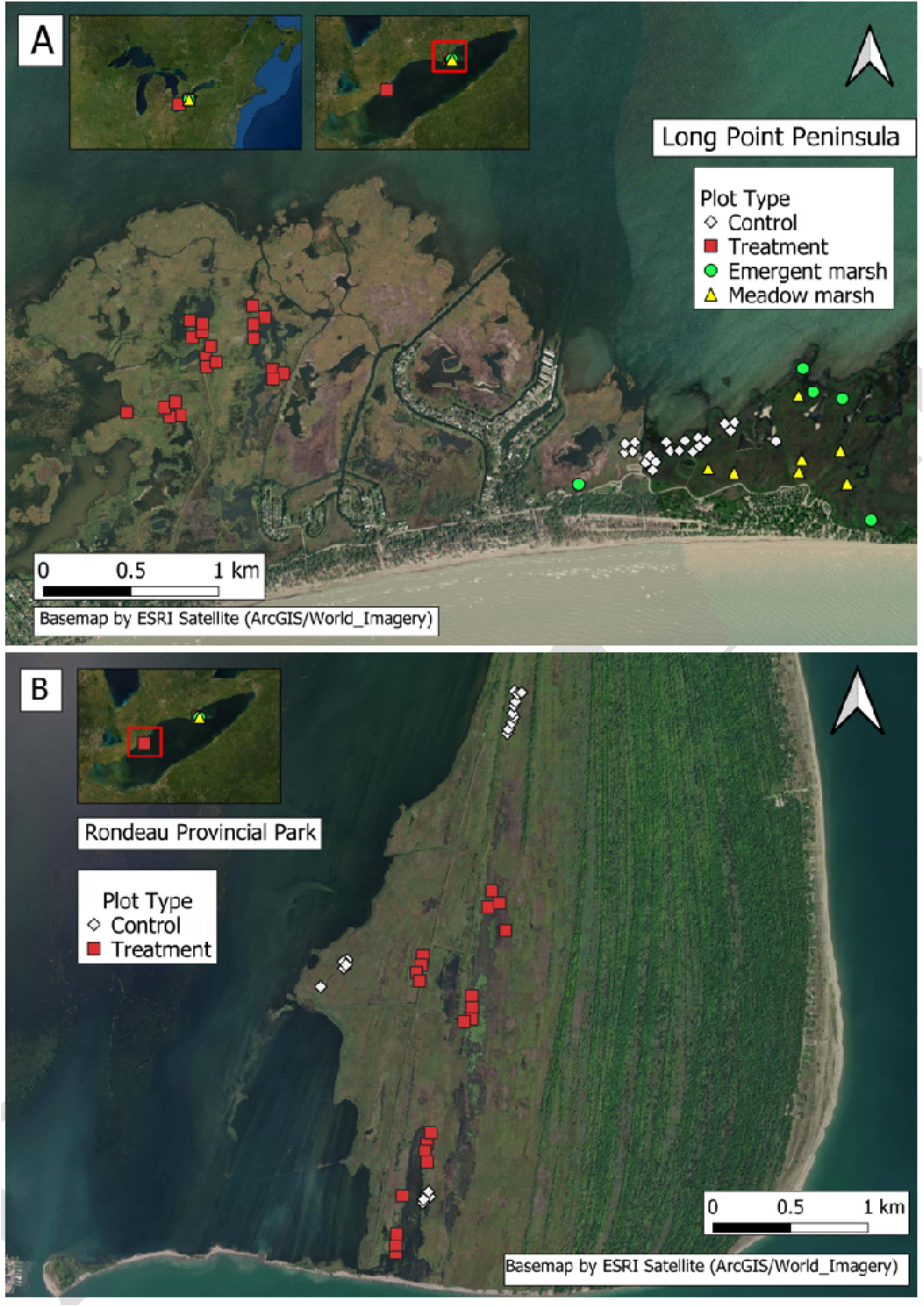
Location of control, treatment and reference plots to assess efficacy of glyphosate-based herbicide treatment on *P. australis* in the western portion of Long Point peninsula (A) and Rondeau Provincial Park (B), located on the north shore of Lake Erie. Reference condition plots were established in Long Point (A) in meadow marsh and emergent marsh communities. Each reference condition point consists of three 1 m^2^ plots spaced a minimum of 10 m apart.

### *Phragmites australis* aerial treatment

The Ontario Ministry of Natural Resources and Forestry (OMNRF), in partnership with the Nature Conservancy of Canada (NCC), and the Ontario Ministry of Environment, Conservation and Parks (OMECP) obtained an Emergency Registration (#32356) under the Pest Control Products Act from Health Canada’s Pest Management Regulation Authority and a provincial Permit to Perform an Aquatic Extermination of invasive *Phragmites australis* in standing water. In September 2016, 335 ha of *P. australis* was treated by licensed contractors in the western portion of Long Point peninsula, and 100 ha was treated in Rondeau Provincial Park. The treatment used Roundup® Custom for Aquatic & Terrestrial Use Liquid Herbicide, combined with a non-ionic alcohol ethoxylate surfactant (Aquasurf®, Registration Number 32152). 4210 g acid equivalent glyphosate/ha as an isopropylamine salt, combined with Aquasurf® non-ionic surfactant at 0.5 L/ha was applied by helicopter (Eurocopter A-Star equipped with GPS guidance and Accu-flo boom nozzles) at a rate of 8.77 L/ha, with a total spray mix of 70 L/ha. In February 2017, the treated marsh in Long Point was mowed and/or rolled to knock down standing dead culms of *P. australis*. Mechanical treatment did not take place in Rondeau Provincial Park.

### Field methods

In August 2016, we established eighty 1 m^2^ plots in Long Point peninsula (n = 40) (Fig. 1A) and Rondeau Provincial Park (n = 40) (Fig. 1B). We then marked plot corners with flagging tape, metal stakes, and a GPS/GNSS unit with sub-meter accuracy (SX Blue II, Geneq Inc., Montreal, PQ, Canada) to ensure the exact locations could be resampled in subsequent years. We situated the plots in dense *P. australis* (>20 stems m^-2^) in a stratified-random manner, such that, in each marsh, treatment (n = 20 per marsh) and control (n = 20 per marsh) plots were paired by water depth, ranging from 10 cm to 48 cm deep (SM1). This represented the range of standing water depths across which dense *P. australis* naturally occurred in our study area (Fig 2). Low density *P. australis* patches were excluded as they were not candidates for herbicide application.

**Figure 2.**
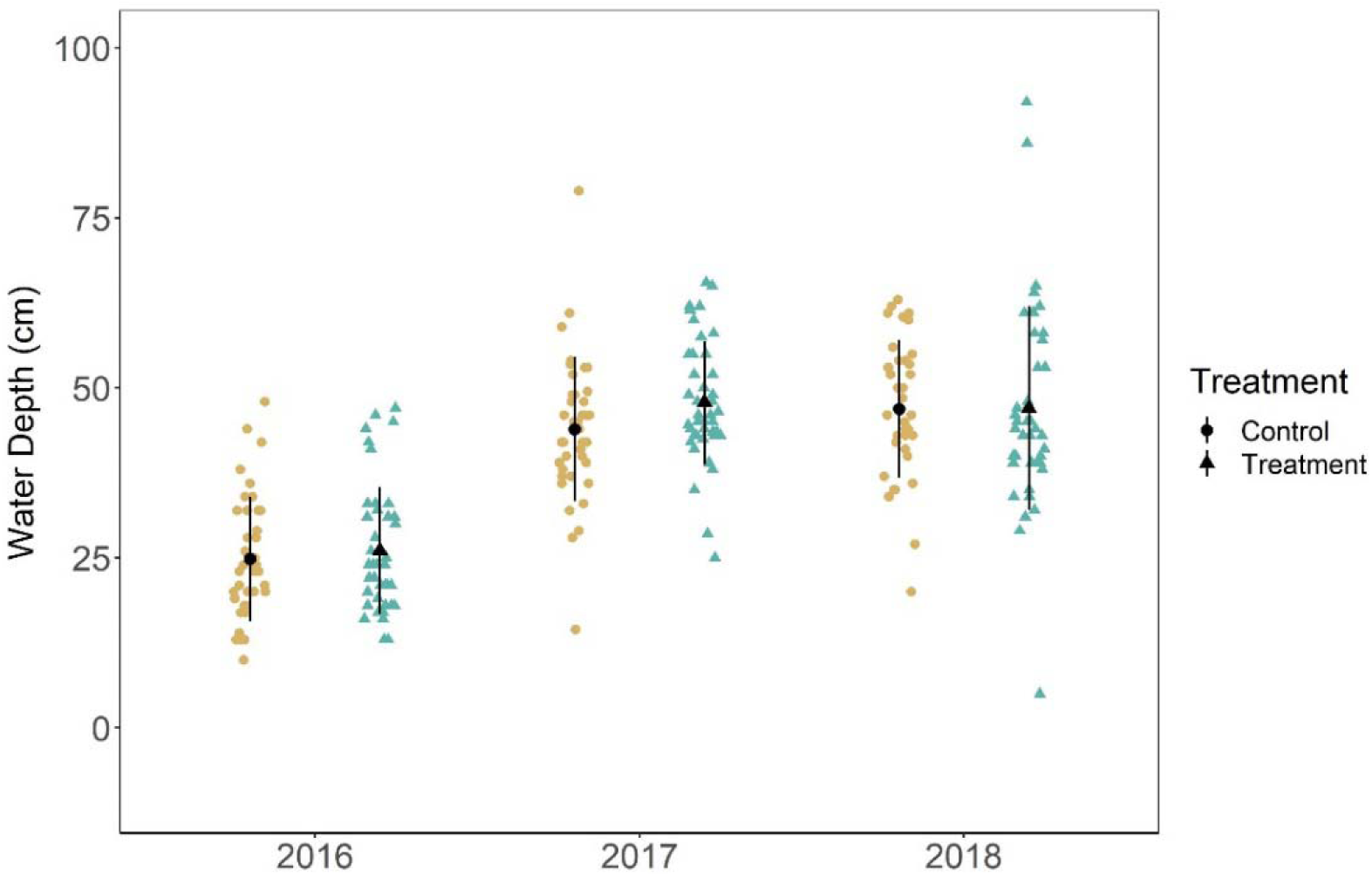
Differences in standing water depth between the control and treatment plots among the three years. Plots in Long Point and Rondeau Provincial Park are combined. Figure created with ggplot2 (Whickham 2016).

All plots were surveyed in August 2016, before treatment, and re-surveyed in August 2017 and August 2018: one- and two-years post-herbicide application, respectively. Importantly, in Rondeau Provincial Park, one control plot was accidentally sprayed with herbicide. Thus, this plot was re-coded as a treated plot, resulting in 39 control and 41 treatment plots. In 2018, a second Rondeau Provincial Park control plot became inaccessible, leaving 38 control and 41 treatment plots. During surveys, we measured relevant ecological variables and vegetation community composition. We characterized the vegetation community composition of the plots based on the percent cover of all plant species and non-living cover, including litter, standing dead, and open water, using a modified Braun-Blanquet cover-abundance method (Wikum and Shanholtzer 1978). Percent cover was considered from a single canopy layer so that each quadrat added up to 100% (± 10%), and species present at less than 1% cover were recorded as 0.05% to document their presence. All *P. australis* stems, living and dead, were counted in each quadrat. Percent incident light reaching the substrate or water surface was measured using a LI-COR LI-1500 light sensor logger with two LI-190 Quantum sensors that measure photosynthetically active radiation (PAR) in the 400 to 700 nm waveband in µmol s^-1^ m^-2^. This permitted simultaneous measurement of the intensity of incident PAR and PAR passing through the canopy. As with any long-term monitoring study, precautions were implemented to reduce damage to the plots such as avoiding trampling, taking care during surveys, and collecting voucher specimens for identification from outside the plot.

In 2017 and 2018 in Long Point, we also characterized the resident vegetation community composition (henceforth “reference condition”) along a similar range of water depths (10 - 56 cm) which encompassed meadow marsh (shallow standing water, hummock forming sedges (*Carex* spp.) and grasses (e.g. *Calamagrostis canadensis* ((Michx.) P. Beauv) and emergent marsh (deeper standing water, robust emergent vegetation (e.g. *Typha* spp.)). We established thirty 1 m^2^ plots in 2017, and twenty-five in 2018, with all plots a minimum of 10 m apart. As the lowest water depth in the control and herbicide-treated plots were 10 cm when established, we removed any reference condition plot with a water depth lower than 10 cm (none in 2017, four in 2018) for analysis. As such there were a total of thirty plots in 2017 (meadow: n = 15, emergent: n = 15), and 21 plots in 2018 (meadow: n = 9, emergent: n = 12).

### Statistical Methods

#### *Efficacy of* P. australis *suppression*

As a significant interaction effect is the hallmark of an effective treatment in a BACI design, we used two-way ANOVAs (type III SS) with treatment (control or herbicide-treated) and year (2016 - 2018) as fixed factors to test for the effect of herbicide application on total and live *P. australis* stem density. We also applied two-way ANOVAs (type III SS) to test for differences in canopy height (cm), and percent incident light (%). To meet assumptions of normality in residuals, we log_10_ transformed percent incident light. As Long Point had secondary treatment to reduce standing dead biomass but Rondeau did not, we also compared total stem density and percent incident light (PAR penetration) in the treatment plots between locations and years (2017 & 2018) with two-way ANOVAs. Where there was a significant effect of a fixed factor, but not the interaction term, we conducted a Tukey’s HSD test. All univariate analyses were carried out using the *car* package (Fox and Weisberg 2019) and *agricolae* package (de Mendiburu 2020) in R v. 3.6.2 (R Core Team 2016).

#### Efficacy of herbicide along a water depth gradient

To assess how effective glyphosate-based herbicide application was along the water depth gradient we compared live *P. australis* stem density in treatment and control plots one year after treatment (i.e., 2017 data) using an ANCOVA, with treatment as a fixed factor and water-depth as a covariate.

### Recovery of Native Vegetation

#### Vegetation community response to treatment

To assess if the vegetation community composition changed in response to herbicide treatment, we conducted a two-way perMANOVA with treatment and year (2016 – 2018) as fixed factors. We applied a general relativization so that percent cover added up to 100% for each plot and removed any species that occurred in two or fewer plots (10 species were removed). Because the number of control and treatment plots was unequal, we used random sampling with replacement that was stratified based on treatment and year. Thirty-eight plots from each treatment and year combination were randomly chosen for each perMANOVA iteration, which we ran 500 times, and we then took the average of each test statistic. This was performed using PC-ORD 7 (McCune and Mefford 2015).

#### Comparing herbicide-treated plots to a reference condition

To visualize vegetation composition changes in the treatment plots from 2016 to 2018, and assess their trajectory towards the *P. australis* control plots or the reference condition plots, we conducted a non-metric multidimensional scaling (NMDS) ordination using a Bray-Curtis dissimilarity matrix. We combined the vegetation composition data from the control and herbicide-treatment plots with the data from the reference condition plots, which we coded as either meadow marsh or emergent marsh based on water depth and dominant vegetation. We then conducted a general relativization so each plot summed to 100%, and removed any species that had two or fewer occurrences in the dataset (17 total). This was performed using PC-ORD 7 (McCune and Mefford 2015).

We then determined the movement of the treatment plots in species space relative to the control and reference condition plots. We defined three centroids (control plots, meadow marsh reference plots, and emergent marsh reference plots) by averaging the NMDS coordinates across all the years for each plot type, and then calculated the Euclidean distance between treatment plots and the three centroids each year. To assess the differences in Euclidean distances among the years we conducted three one-way ANOVAs with “distance to centroid” and “year” as fixed factors. If the model was significant, we conducted a Tukey’s HSD post-hoc test. Analyses were conducted using the *car* (Fox and Weisberg 2019) and *agricolae* (de Mendiburu 2020) packages in R v. 3.6.2 (R Core Team 2016). In addition to assessing the average movement of treatment plots relative to the three centroids each year, we identified the trajectory of individual treatment plots. We assigned each plot a ‘reference condition’ based on the reference centroid it was closest to in 2018, and calculated the total distance between the plot, its reference centroid, and the control centroid in both 2017 and 2018. We then calculated the distance between the plot and reference condition as a proportion of the total distance, such that a smaller value indicates that the plot is closer to its reference condition.

## Results

### *Suppression of* P. australis *regrowth*

For every variable related to *P. australis* suppression there was a significant interaction between treatment and year, indicating that herbicide successfully suppressed *P. australis* in treated plots (Fig. 3; SM2). In 2017, we observed a 99.7% reduction in live *P. australis* stem density in treated plots compared to control plots: on average there were 0.1 (std. = 0.6) live *P. australis* stems/m^2^ in treatment plots compared to 36.8 (std. = 15.3) live stems/m^2^ in control plots (SM3). All the live *P. australis* stems (n = 4) were concentrated in a single treated plot. In 2018, there was a 94.7% decrease in live *P. australis* stem density in treated plots compared to control plots: an average of 1.5 (std. = 5.6) live stems/m^2^ in treatment plots and 29.8 (std. = 12.3) live stems/m^2^ in control plots (Fig. 3A). Specifically, live *P. australis* was concentrated among 5 plots and summed to 62 live stems.

**Figure 3.**
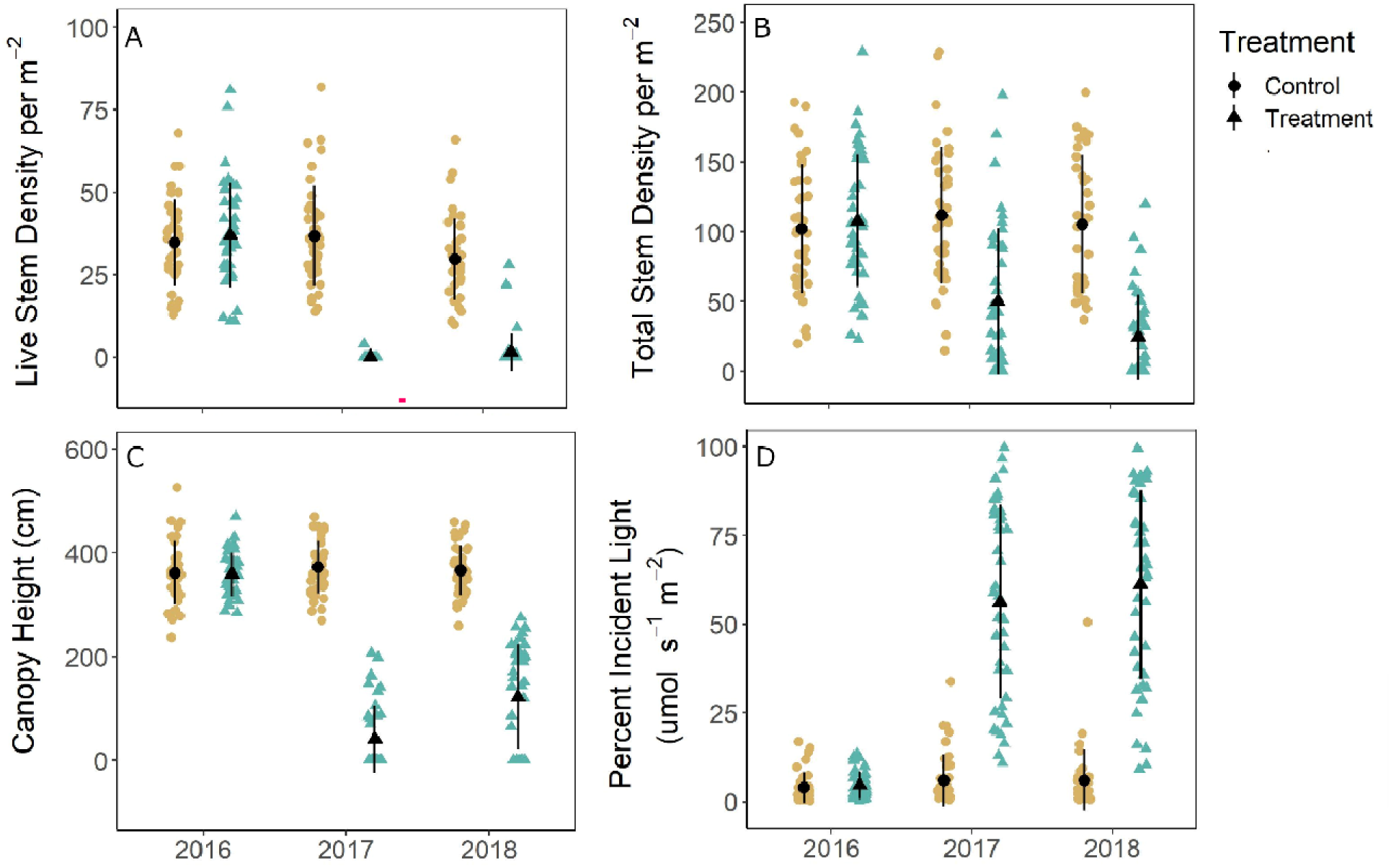
There was a significant interaction between treatment type and year for all variables related to *P. australis* suppression: live *P. australis* stems/m^2^ (A), total *P. australis* stems/m^2^ (B), canopy height (cm) (C), and percent incident light reaching substrate (D). This represents the clear suppression of *P. australis* glyphosate-based herbicide application to targeted areas. Error bars represent the standard deviation. Created with ggplot2 (Whickham 2016).

There was no difference in live stem density between Long Point and Rondeau Provincial Park after treatment (two-way ANOVA F_1,78_ ≤ 0.883, p ≥ 0.350; SM4). However, there was a significant interaction between year and location for total stem density (two-way ANOVA F_1,78_ = 4.286, p = 0.042; SM4). In Long Point, where rolling and mowing occurred, there were fewer total (i.e., live plus dead) *P. australis* stems in 2017 (average 14.0 stems/m^2^, std. = 21.5) compared to Rondeau (average 87.6 total stems/m^2^, std. = 49.1). By 2018, there were on average 2.8 total *P. australis* stems/m^2^ (std. = 7.16) in Long Point compared to 44.8 total *P. australis* stems/m^2^ (std. = 30.9) in Rondeau. The percent of PAR reaching the sediment also exhibited a significant interaction between location and year (two-way ANOVA F_1,78_ = 8.460, p = 0.005; SM4). One year after treatment, 68.8% (std. = 26.2%) of PAR reached the substrate in Long Point compared to 44.3% (std. = 23.2%) in Rondeau. However, in 2018 values were similar in both marshes, with an average of 56.2% (std. = 28.1%) in Long Point and 66.8% (std. = 25.0%) in Rondeau.

### Efficacy of herbicide along a water depth gradient

There was no interaction between the effect of treatment and water depth on the density of live *P. australis* in the year after treatment (ANCOVA F_1,77_ = 0.043, p = 0.836). Herbicide-treatment had a significant effect on the density of live *P. australis* stems (ANCOVA F_1,77_ = 988.50, p < 0.001), while water depth did not (ANCOVA F_1,77_ = 0.076, p = 0.784; SM5) indicating glyphosate-based herbicide worked equivalently at all water depths.

### Recovery of Native Vegetation

#### Vegetation community BACI response

The vegetation community composition in the plots that where treated with a glyphosate-based herbicide was compositionally different from the control plots following herbicide treatment (perMANOVA, pseudo-F_2, 222_ = 44.77 (std. = 5.33), p = 0.001 (std. < 0.001); SM6). In 2016, we observed a total of 26 species of vascular plants in control plots and 31 species in treatment plots (Fig. 5). *Phragmites australis* was the most abundant species in both treatments, with an average of 73.8% cover (std. = 13.3%) in control plots and 73.6% (std. = 15.0%) in treatment plots. In 2017, we identified 26 species in control plots, and again the most abundant species was *P. australis* (66.7% std. = 17.5%). In the treatment plots we identified 22 species and the most abundant species was *Hydrocharis morsus-ranae* (L.) (European frogbit) (33.6% cover std. = 29.8%), a small floating aquatic plant. In 2018, we identified 17 species in control plots, with *P. australis* the most abundant (85.7%, std. = 12.8%) and 21 species in treatment plots, where *H. morsus-ranae* was the most abundant (48%, std. = 38.7%). Average species richness per plot decreased slightly each year, although there were no significant differences between treatments (two-way ANOVA F_1,233_ = 0.369, p = 0.544), among years (two-way ANOVA F_2,233_ = 3.005, p = 0.051), nor an interaction (two-way ANOVA F_2,233_ = 2.869, p = 0.059).

**Figure 4.**
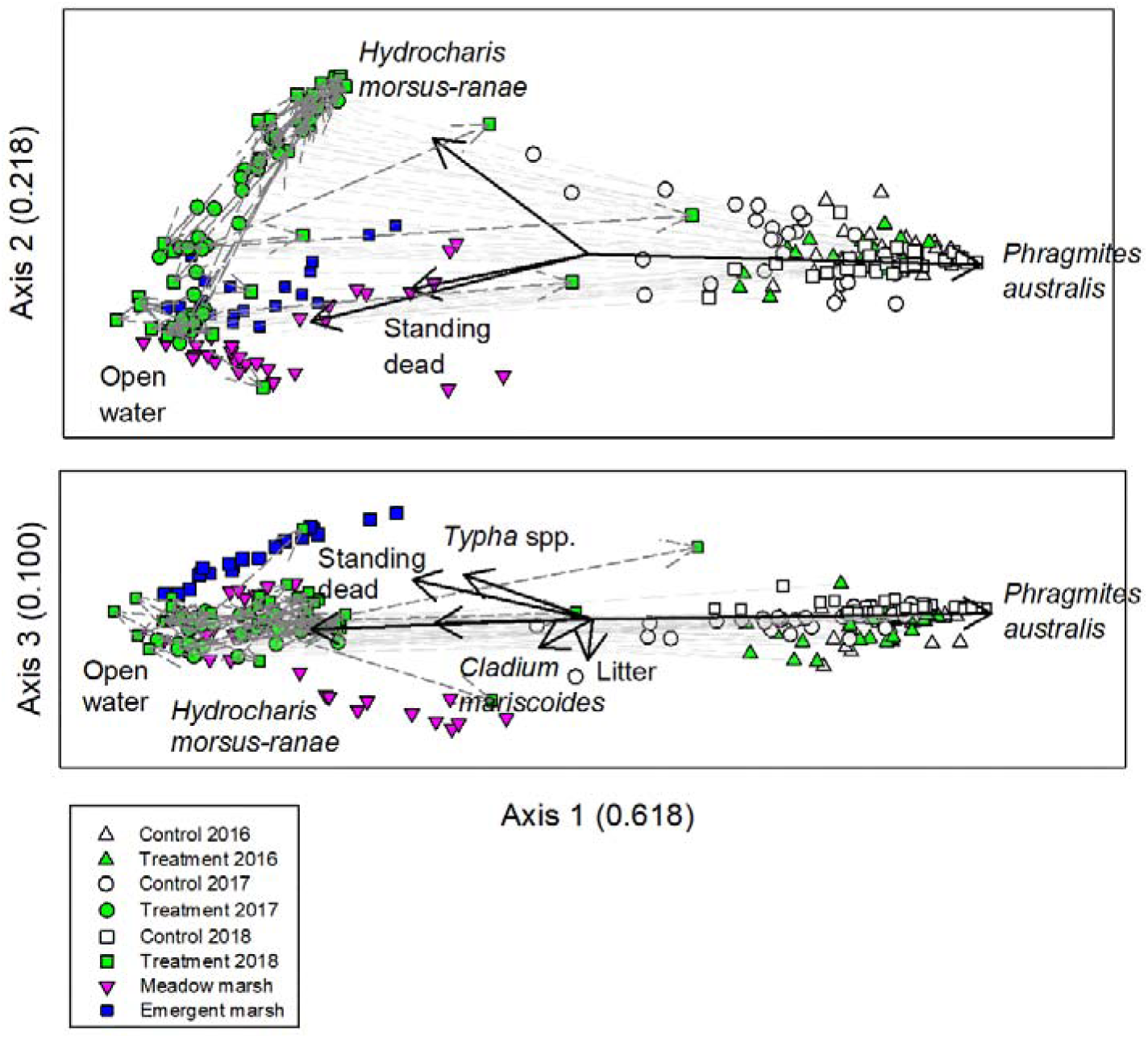
The final 3D NMDS ordination solution, using Bray-Curtis dissimilarity matrix. Control and treatment plots were measured in 2016, before glyphosate-based herbicide treatment occurred, and in 2017 and 2018. Reference plots were sampled in 2017 and 2018. Black vectors represent reasonably correlated (r^2^ ≥ 0.150) cover classes. Light gray vectors represent the movement of treatment plots between 2016 and 2017, and dark gray vectors represent the movement of treatment plots between 2017 and 2018. Note axes are scaled by the proportion of variance in community composition they explained.

**Figure 5.**
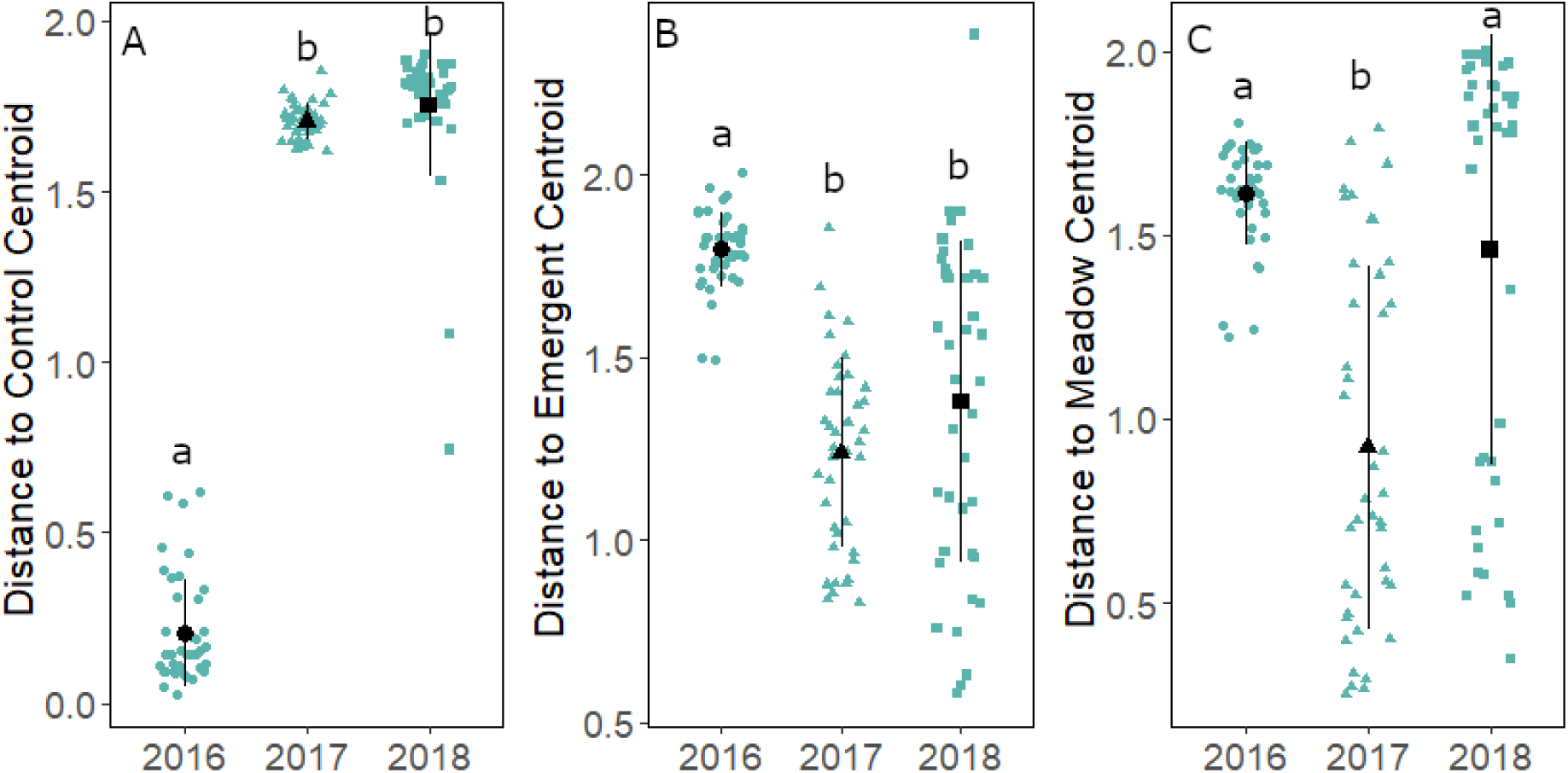
Euclidean distances between the treatment plots (n = 41) and the control plot centroid (A), the emergent marsh plot centroid (B), and the meadow marsh plot centroid (C) in terms of their vegetation community composition. The smaller the distance, the more closely the vegetation community in that treatment plot resembles the vegetation community typical of the specified plot type. Error bars represent the standard deviation, lower case letters represent significant differences based on Tukey’s HSD. Created with ggplot2 (Whickham 2016).

#### Comparing treated plots to a reference condition

Our final NMDS ordination depicts the differences in community composition between treatments among the years and compares community composition in treated plots to reference condition plots (Figure 4). The final NMDS ordination was a 3-dimensional solution, with an instability of < 0.0001 after 121 iterations, and a stress of 8.94. The proportion of total variance explained was 0.931 (axis 1: 0.618, axis 2: 0.213, axis 3: 0.100). All species and cover class vectors and their correlations with site scores are presented in SM7. We also conducted an NMDS ordination containing only the control and treatment plots (SM8 & SM9) that exhibits the same general patterns as displayed in figure 5.

The differences along NMDS axis 1 are driven primarily by *P. australis* abundance. In 2016, both control and treatment plots cluster in an area of high *P. australis* abundance whereas in 2017 the treatment plots create a new space in the ordination defined by a vegetation community that contains no *P. australis*. Of note, the meadow and emergent marsh plots cluster with the treatment plots rather than creating their own space. This is promising, as it means that the herbicide-treated plots became more similar to reference condition communities after treatment. Axis 2 illustrates a gradient of open-water and floating vegetation extent. Reference condition plots have more open-water, indicating higher structural heterogeneity, whereas many treatment plots are dominated by *H. morsus-ranae*. In 2018, the treatment plots begin to form two clusters along this gradient – those plots with a high abundance of *H. morsus-ranae*, and those that are more similar to the reference condition plots. Many treatment plots exhibited an increase in *H. morsus-ranae:* before treatment, the average percent cover was 1.73% (std = 2.86%), one-year after treatment it was 33.6% (std = 29.8%) and two-years afterwards it was 48.0% (std = 38.7%).

Finally, axis 3 separates the meadow marsh and emergent marsh communities that comprise the reference condition plots. Meadow marsh was more diverse than emergent marsh; we identified 23 species (7.9 per plot std. = 2.5) in 2017 and 22 species (8.2 per plot std = 2.9) in 2018. The meadow marsh community is characterized by *Calamagrostis canadensis* (Canada Blue-joint grass), a mix of sedges (e.g., *Carex aquatilis* (Wahlenb.) and *Carex lasiocarpa* (Ehrh.)) and other herbaceous vegetation (e.g.,, *Cladium mariscoides* ((Muhl.) Torr.) (Smooth Sawgrass)). In contrast, in emergent marsh we identified 8 species (4.0 per plot std = 1.0) in 2017 and 6 species (4.2 per plot std = 1.2) in 2018. The emergent marsh community is characterized by high *Typha* spp. abundance, likely the majority is hybrid *Typha* x *glauca* (Godr. (pro sp.)) though field identification based on morphology can be challenging. On the third axis, the treated plots fall in the middle of the two reference communities, indicating they are more similar to reference than control plots, but do not exhibit the same composition as either reference community, even two years after treatment.

The assessment of Euclidean distances demonstrates the trajectory that vegetation in the treatment plots followed over time, allowing us to interpolate between the control condition on the one hand (Fig. 5A), and the reference conditions on the other (Fig. 5B, C). Prior to herbicide application in 2016, all treatment plots closely resembled control plots, but in 2017 the vegetation community in treated plots differed substantially from control plots (one-way ANOVA F_2, 120_ = 1348.00, p < 0.001; Fig. 5A). In 2018, the treated plots that had higher re-growth of *P. australis* moved back toward the control plot centroid (Fig. 5A).

In 2017, nearly all plots increased in similarity to the emergent marsh (one-way ANOVA F_2, 120_ = 37.911, p < 0.001; Fig. 5B) and meadow marsh (one-way ANOVA F_2, 120_ = 26.741, p < 0.001; Fig. 5C) reference condition centroids. In 2018, rather than follow a continuous trajectory toward the reference conditions, a subset of treated plots increased in distance from the two reference conditions, particularly the meadow marsh condition (Fig. 5C). This is not matched by a reduction in Euclidean distance between the plots and the control centroid (Fig. 5A), indicating this was not attributable to regrowth by *P. australis*. Instead, these treatment plots moved into novel species space, undefined by the interpolation between control and reference conditions. The vegetation community in only 16 of the 41 treatment plots more closely resembled the best fitting reference condition in 2018 than they did in 2017 (SM10). Of these, 10 were most similar to emergent marsh and 6 were most similar to meadow marsh habitats. Two treatment plots with a high abundance of *P. australis* became more similar to the control plots by 2018.

## Discussion

This project represents the first large-scale application (455 ha treated in 2016) of a glyphosate-based herbicide over water to control invasive *P. australis* in Canada. We assessed the efficacy of this unprecedented management action at suppressing *P. australis* populations and the recovery of native vegetation communities in two marsh complexes on the north shore of Lake Erie: Long Point peninsula and Rondeau Provincial Park.

### *Suppression of* P. australis

The suppression efficacy reported by studies of glyphosate-based *P. australis* management varies quite substantially from lows of 50-60% (e.g., Farnsworth and Meyerson 1999; Ailstock et al. 2001) to highs of >90% (e.g., Derr 2008; Zimmerman et al. 2018). A comparison of marsh types, plot sizes, and inclusion of secondary treatments like mowing, mulching or burning post-herbicide application does not explain the differences among reported suppression efficacy in the literature. The percent reduction in *P. australis* stem density in our study is on the high end of the range reported in similar studies in marshes. Glyphosate-based herbicide was very effective at suppressing *P. australis* for at least two years after treatment, with a 99.7% reduction in live stem density per plot in the first year and a 94.7% reduction in the second year, relative to control plots. In the first year following treatment, *P. australis* remained in only one of 40 treated plots (97.5% reduction). By the second year, *P. australis* had recolonized four additional plots, such that 87.5% of treated plots remained free of *P. australis*.

We found no effect of water depth on the efficacy of *P. australis* suppression. The glyphosate-based herbicide was equally as effective across the water depth gradient along which dense *P. australis* naturally occurred (10 – 48 cm). In semiarid regions, drier sites may result in less successful herbicide-based *P. australis* suppression, as water stress limits the translocation of the herbicide in the plant (Rohal et al. 2019a). Our results suggest that, in an area with adequate moisture for *P. australis*, the actual water depth does not affect how well the herbicide works as water depth does not inhibit adsorption by plant leaves and translocation into rhizomes.

### *Re-growth of* P. australis

We observed minimal *P. australis* re-growth during the two years after herbicide treatment. Inundation reduces the likelihood that *P. australis* will germinate from seeds (Armstrong et al. 1999), therefore we conclude that the majority of the re-growth in our study was the result of surviving rhizomes, as water depths ranged from 13 cm to 92 cm, with the exception of one site, over the three years of our study. The first year after treatment, only one of forty treated plots (2.5%) had live *P. australis*. It contained four live ramets in the first year, but expanded seven-fold to 28 ramets in the subsequent year, which is equivalent to the average of 29.8 (std. = 12.3) live stems/m^2^ in control plots in 2018. By the second year after treatment four more plots were re-invaded, at densities from 1 to 22 live *P. australis* stem/m^2^. This rapid expansion and re-invasion from small remnants of *P. australis* mirrors the results of long-term monitoring studies (e.g., Lombard et al. 2012; Quirion et al. 2018). For example, after glyphosate was used to suppress *P. australis* in a forest preserve in upstate New York, 13.6% of sites exhibited re-growth the following year (Quirion et al. 2018). Since *P. australis* can rapidly overtake an area from small remnant patches, it is important to recognize that *P. australis* management entails two distinct phases. Initial large-scale treatment with herbicide reduces the extent and density of established *P. australis*, achieving the objective of broad ‘suppression’. Afterwards, management enters a second ‘containment’ stage, where follow-up spot treatment is required as part of routine maintenance. Quirion et al. (2017) determined continued containment and spot treatment can keep maintenance costs low, as the probability of re-invasion significantly decreased as the *P. australis*-free duration increased, with no reappearance documented after four years of consecutive absence (Quirion et al. 2017). This emphasizes the importance of long-term monitoring and appropriate project budgeting to support the containment phase and to prevent re-establishment of *P. australis* patches.

### Recovery of native vegetation

While herbicide treatment was effective at suppressing *P. australis*, we did not observe an increase in species richness in treated plots, despite reductions in canopy height and increases in PAR penetration. In fact, there was a slight reduction in treatment plot richness compared to control plots, and average species richness declined in both control and treatment plots over the three years. We attribute this to prolonged flooding, as Lake Erie’s water levels have been high since 2016 (DFO 2019). Prolonged high water can negatively impact marsh vegetation (Van Der Valk 2005), and recent work by Keddy and Campbell (2019) suggests that four consecutive years of flooding is enough to drown marsh plants in Lake Erie coastal marshes. Prolonged flooding also limits seedbank regeneration (Keddy and Reznicek 1986). Meadow marsh in particular is less flood tolerant than other vegetation communities, and is typically among the most species-rich community in coastal marshes (Reznicek and Catling 1989; Keddy and Campbell 2019). Yet maintaining floooding over 30 cm kills *P. australis* seedlings and slows or stops stand expansion (Norris et al. 2002). As such, it is likely that the high Lake Erie water levels after initial herbicide treatment have aided the suppression of *P. australis*, while limiting native vegetation recovery.

After two years, more than half of the treatment plots had a community composition that was dissimilar to the reference condition plots and the control plots. These plots have a novel vegetation community characterized by floating and submerged aquatic vegetation, with a high abundance of non-native *Hydrocharis morsus-ranae* (European frogbit). *Hydrocharis morsus-ranae* is a small, free-floating aquatic plant native to Europe, Asia, and Africa which was discovered in Canada in 1932 (Catling et al. 2003). *Hydrocharis morsus-ranae* was present in both Long Point and Rondeau before treatment and often co-occurred with *P. australis*. In fact, robust emergent vegetation (i.e. *Typha* spp.) has been demonstrated to facilitate the establishment of *H. morsus-ranae* in Great Lakes wetlands by reducing wave action and wind energy (Monks et al. 2019). A recent review of *H. morsus-ranae* in North America by Zhu et al. (2018) concluded that the dense mats created by *H. morsus-ranae* negatively impact native aquatic plant diversity, while effects on macroinvertebrates vary depending on the invaded system. One study found a negative relationship between *H. morsus-ranae* and benthic invertebrate richness and abundance, though Chironomidae abundance was higher in invaded areas (Zhu et al. 2015). The authors conclude that *H. morsus-ranae* may have fewer negative impacts than anticipated in North American wetlands (Zhu et al. 2018), though more studies on invertebrates and wetland food webs are warranted before we conclude it is less harmful to wetland flora and fauna than the invasive *P. australis* it is replacing.

Secondary invasions by non-native species present a challenge in restoration. Non-native species are common in restored wetlands, and when coupled with a lack of native propagules, can be a reason restoration projects do not meet their targets (Matthews and Spyreas 2010; Bonello and Judd 2020). The suppression of a dominant invasive species often leads to an increase in one or more non-target invasive species with little increase in native species (Pearson et al. 2016). We are not the first to report a secondary invasion following *P. australis* suppression; herbicide-treatment of *P. australis* invaded Great Lakes coastal marshes in Detroit also resulted in dense populations of *H. morsus-ranae* (Judd and Francoeur 2019).

Pearson et al. (2016) suggest secondary invasion is driven by four factors: treatment side effects, target invader legacy effects, provenance effects, and shifting environmental conditions. We expect there are minimal treatment side effects in our system, as aerial application does not compact sediment and overspray was restricted to within 15 m of target treatment areas based on UAV imagery (Adam Hogg, Ontario Ministry of Natural Resources and Forestry *pers comm* 26 March 2018). Further, concentrations of glyphosate did not approach thresholds of toxicological concern and dissipated within 30 days to 1 year from water and sediment, respectively (Robichaud and Rooney, in review). However, legacy effects created by established *P. australis*, such as alterations to soil nutrients (Yuckin and Rooney 2019, D’Antonio and Meyerson 2002), and shifting environmental conditions in and surrounding Lake Erie, for example nutrient pollution (e.g., Mohamed et al. 2019) and climate change (e.g., Zhang et al. 2020), may increase the likelihood of secondary invasions.

The suppression of *P. australis* alone was not sufficient to put treatment plots on a trajectory towards one of the two historically dominant vegetation communities. Wetlands dominated by *P. australis* may represent a degraded stable state, and *P. australis* suppression has the potential to put the ecosystem on an unintended trajectory (sensu Suding et al. 2004), which in this instance was characterized by invasive *H. morsus-ranae*. The question remains whether, after lake levels drop, native species will be capable of recolonizing and persisting in areas where *P. australis* has been dominant for over two decades. The species present in seedbanks are an important determinant of the success of passive recovery and seedbanks in invaded areas typically are less dense and less species rich (Gioria et al. 2014). Shifting environmental conditions, whether driven by anthropogenic disturbance or climate change, are likely to result in novel ecosystems and alternative stable states (Harris and Hobbs 2006). In our study system, there are numerous factors that lead to the persistence of a degraded state dominated by *H. morsus-ranae*. It is therefore unlikely, given these factors, that suppression of *P. australis* alone will re-create historical vegetation communities. However, this does not mean that suppression is not a worthwhile endeavor. There are already demonstrable benefits to suppressing *P. australis* in these marshes, including an increase in marsh birds (Tozer and Mackenzie 2019) and at-risk plant species (Polowyk 2020), with little associated risk from herbicide-treatment (Robichaud and Rooney, in review). Global and local changes require conservation practitioners to evaluate restoration goals, and acknowledge that in a rapidly changing world some ecosystems may not retain their original composition or function (Hobbs et al. 2009).

### Recommendations

With the current distribution and level of establishment of *P. australis* in North America, most control projects must focus on suppression, containment, and asset protection rather than aim for complete eradication. As the size of invaded area increases, it becomes increasingly difficult to achieve eradication (Quirion et al. 2017) or to recover native vegetation communities (Rohal et al. 2019b). The speed at which *P. australis* begins to re-colonize a treated area means that long-term monitoring and follow-up treatment are essential components of any control project. While eradication may not be possible, substantial ecological benefits can be achieved through continuous maintenance and containment at relatively low costs. For example, Meyer et al. (2010) found high levels of bird use in low density *P. australis*, especially along patch edges. On-going monitoring and follow-up treatment require long term budget allocations to invasive species management, but given that 25% of Ontario’s species at risk are threatened by *P. australis* invasion (Bickerton 2015), we argue that it is a worthwhile investment.

Successful invasive species treatment requires both a reduction of the target population and the recovery of native vegetation. Hence, long-term monitoring is also required to assess the trajectory of the recovering vegetation community. Passive restoration relies on propagules that are present within, or can easily reach, the target area. The diversity in seedbanks where *P. australis* has invaded varies depending on the system and history of invasion, though some studies suggest that even in marsh invaded by *P. australis* for years the seedbank diversity is sufficient for passive restoration (e.g., Howell 2017; Hazelton et al. 2018). More, surveys of the seedbank can inform managers about the risks of secondary invasion and the capacity of the seedbank to support native species recolonization of treated areas. Based on the high likelihood of secondary invasion, those undertaking invasive species control projects should identify underlying causes that contribute to degradation (Gaertner et al. 2012) as well as treatment side effects and invader legacies that make sites more susceptible to secondary invasion (Pearson et al. 2016).

## Declarations

### Code availability

A GitHub repository of all relevant code can be made public upon acceptance

## Authors’ contributions

Robichaud conducted field work, statistical analyses, and prepared the manuscript. Rooney conceived of and designed the study, developed the field protocols, coordinated the field team, and assisted with manuscript preparation.

## Acknowledgements

We would like to thank the graduate students and technicians who made this project possible: Graham Howell, Sarah Yuckin, Daina Anderson, Jody Daniel, Heather Polowyk, and Calvin Lei. Thank you to the Ontario Ministry of Natural Resources and Forestry, the Nature Conservancy of Canada, and Expedition Helicopters of Cochrane who carried out the aerial application of herbicide.

