## Supplemental materials 1-8 for "Suppression of established invasive *Phragmites australis* leads to secondary invasion"

SM1. Water depths from every control and treatment plot in Long Point (LP) and Rondeau Provincial Park (RPP). Control (n = 39) and treatment (n = 41) plots were paired by August 2016 water depths across the range occupied by *P. australis* at sufficient density to warrant herbicide application (> 20 stem $m^{-2}$). Control plot RCP16 was mistakenly treated and thus was re-coded.  Further, control plot RPC41 became inaccessible in 2018, reducing the number of control plots to 38.

| SiteID | Location | Treatment | Water Depth (cm) | | |
|---|---|---|---|---|---|
| | | | 2016 | 2017 | 2018 |
| LPC25 | LP | Control | 10 | 38 | 27 |
| LPC24 | LP | Control | 13 | 36 | 44 |
| LPC39 | LP | Control | 13 | 28 | 20 |
| RPC04 | RPP | Control | 13 | 29 | 37 |
| RPC15 | RPP | Control | 13 | 14.5 | 34 |
| LPC26 | LP | Control | 14 | 39 | 52 |
| LPC27 | LP | Control | 17 | 42 | 50 |
| RPC14 | RPP | Control | 17 | 36 | 56 |
| LPC33 | LP | Control | 18 | 32 | 34 |
| LPC37 | LP | Control | 18 | 46 | 40 |
| LPC32 | LP | Control | 19 | 40 | 46 |
| LPC31 | LP | Control | 20 | 46 | 36 |
| RPC05 | RPP | Control | 20 | 37 | 50 |
| RPC08 | RPP | Control | 20 | 45 | 46 |
| RPC18 | RPP | Control | 20 | 39 | 53 |
| LPC21 | LP | Control | 21 | 52 | 60 |
| RPC12 | RPP | Control | 21 | 42 | 50 |
| LPC28 | LP | Control | 23 | 44 | 54 |
| RPC11 | RPP | Control | 23 | 46 | 43 |
| RPC20 | RPP | Control | 23 | 40 | 43 |
| LPC23 | LP | Control | 24 | 59 | 52 |
| LPC36 | LP | Control | 24 | 45 | 45 |
| RPC10 | RPP | Control | 24 | 41 | 48.5 |
| RPC07 | RPP | Control | 25 | 46 | 53.5 |
| LPC30 | LP | Control | 26 | 49 | 41 |
| LPC35 | LP | Control | 28 | 54 | 43 |
| LPC40 | LP | Control | 28 | 33 | 35 |
| LPC29 | LP | Control | 29 | 53 | 61 |
| RPC03 | RPP | Control | 32 | 49 | 60.5 |
| RPC06 | RPP | Control | 32 | 42 | 62 |
| RPC09 | RPP | Control | 32 | 37 | 53 |
| RPC17 | RPP | Control | 32 | 49.5 | 61 |
| LPC38 | LP | Control | 34 | 53 | 35 |
| RPC13 | RPP | Control | 34 | 42 | 42 |
| LPC22 | LP | Control | 36 | 48 | 63 |
| LPC34 | LP | Control | 38 | 61 | 44 |
| RPC02 | RPP | Control | 42 | 53.5 | 54 |

| | | | | | |
|---|---|---|---|---|---|
| RPC19 | RPP | Control | 44 | 48 | 55 |
| RPC41 | RPP | Control | 48 | 79 | NC |
| LPT13 | LP | Treatment | 13 | 43 | 39 |
| LPT41 | LP | Treatment | 13 | 43 | 32 |
| RPT28 | RPP | Treatment | 16 | 35 | 39 |
| RPT34 | RPP | Treatment | 16 | 28.5 | 45 |
| LPT15 | LP | Treatment | 17 | 44.5 | 35 |
| RPT32 | RPP | Treatment | 17 | 39 | 34 |
| LPT12 | LP | Treatment | 18 | 46.5 | 39 |
| LPT19 | LP | Treatment | 18 | 52 | 62 |
| RPT22 | RPP | Treatment | 18 | 52 | 53 |
| RPT37 | RPP | Treatment | 18 | 41 | 47 |
| LPT16 | LP | Treatment | 19 | 42 | 31 |
| RPT31 | RPP | Treatment | 20 | 38 | 45 |
| LPT08 | LP | Treatment | 21 | 45 | 40 |
| LPT18 | LP | Treatment | 21 | 55 | 40 |
| RPT25 | RPP | Treatment | 21 | 45 | 53 |
| RPT39 | RPP | Treatment | 21 | 42.5 | 45 |
| LPT07 | LP | Treatment | 22 | 43.5 | 44 |
| LPT10 | LP | Treatment | 22 | 60 | 41 |
| LPT14 | LP | Treatment | 22 | 43.5 | 43 |
| RPT30 | RPP | Treatment | 22 | 46 | 58 |
| LPT09 | LP | Treatment | 24 | 49 | 34 |
| LPT20 | LP | Treatment | 24 | 43 | 38 |
| LPT42 | LP | Treatment | 24 | 55 | 47 |
| RPT40 | RPP | Treatment | 24 | 44 | 46 |
| LPT43 | LP | Treatment | 25 | 50 | 39 |
| RPT24 | RPP | Treatment | 26 | 49 | 58 |
| LPT11 | LP | Treatment | 28 | 58 | 44 |
| LPT06 | LP | Treatment | 30 | 43 | 43 |
| LPT05 | LP | Treatment | 31 | 65 | 43 |
| RPT23 | RPP | Treatment | 31 | 55 | 61 |
| RPT33 | RPP | Treatment | 31 | 46 | 57 |
| RPT29 | RPP | Treatment | 32 | 45 | 40 |
| LPT44 | LP | Treatment | 33 | 61.5 | 29 |
| RPT21 | RPP | Treatment | 33 | 62 | 64 |
| RPT38 | RPP | Treatment | 33 | 48 | 61 |
| RPC16 | RPP | Treatment | 41 | 25 | 48 |
| LPT17 | LP | Treatment | 42 | 62 | 5 |
| RPT27 | RPP | Treatment | 44 | 57.5 | 65 |
| RPT36 | RPP | Treatment | 45 | 44 | 92 |
| RPT35 | RPP | Treatment | 46 | 48 | 61 |
| RPT26 | RPP | Treatment | 47 | 65.5 | 86 |

SM2. Two-way ANOVA results, type III SS with year (2016, 2017, 2018) and treatment as fixed factors. The percent of PAR penetrating the canopy was $\log_{10}$ transformed. There was a significant interaction for every measured variable.

| | Live stems (per m$^2$) | | | Total stems (per m$^2$) | | | Canopy Height (cm) | | | PAR penetration (% incident light) | | |
|---|---|---|---|---|---|---|---|---|---|---|---|---|
| | df | F | P | df | F | P | df | F | P | df | F | P |
| Year | 2 | 3.615 | 0.028 | 2 | 0.448 | 0.639 | 2 | 0.267 | 0.766 | 2 | 1.697 | 0.186 |
| Treatment | 1 | 0.735 | 0.392 | 1 | 0.312 | 0.577 | 1 | 0.047 | 0.828 | 1 | 1.243 | 0.266 |
| Year x Treatment | 2 | 60.612 | < 0.001 | 2 | 19.293 | < 0.001 | 2 | 138.250 | < 0.001 | 2 | 51.208 | < 0.001 |
| Error | 233 | | | 233 | | | 233 | | | 233 | | |

SM3. The average, standard deviation, minimum and maximum live *P. australis* stem density, total (live & dead) stem density, canopy height and photosynthetically active radiation (PAR) penetration in both treatments over the three years; pre-treatment (2016) and two years post.

| | | Live Stems (per $m^{-2}$) | | | Total Stems (per $m^{-2}$) | | | Canopy Height (cm) | | | PAR penetration (% incident light) | | |
|---|---|---|---|---|---|---|---|---|---|---|---|---|---|
| | | Avg. (std.) | Min. | Max. | Avg. (std.) | Min. | Max. | Avg. (std.) | Min. | Max. | Avg. (std.) | Min. | Max. |
| Control | 2016 | 34.7 (13.1) | 13 | 68 | 102 (46.2) | 20 | 193 | 362 (61.1) | 237 | 526 | 4.1 (4.3) | 0.4 | 17.0 |
| | 2017 | 36.8 (15.3) | 14 | 82 | 112 (49.0) | 15 | 229 | 372 (50.4) | 270 | 470 | 6.0 (7.5) | 0.6 | 33.8 |
| | 2018 | 29.8 (12.3) | 10 | 66 | 105 (49.5) | 37 | 200 | 367 (47.4) | 260 | 460 | 6.0 (8.6) | 0.7 | 50.6 |
| Treatment | 2016 | 37.0 (15.8) | 11 | 81 | 108 (47.5) | 23 | 229 | 358 (41.5) | 285 | 470 | 4.7 (3.9) | 0.3 | 13.6 |
| | 2017 | 0.1 (0.6) | 0 | 4 | 50.2 (52.3) | 0 | 120 | 40.7 (65.5) | 0 | 207 | 56.3 (27.4) | 11.0 | 99.6 |
| | 2018 | 1.5 (5.6) | 0 | 28 | 24.3 (30.5) | 0 | 198 | 121 (102) | 0 | 275 | 61.1 (26.6) | 9.0 | 99.4 |

SM4. Two-way ANOVA results, type III SS, comparing total stem density, live stem density, and photosynthetically active radiation (PAR) penetration in 2017 and 2018 between Long Point and Rondeau Provincial Park. Secondary treatment (e.g. rolling and mowing) occurred in Long Point but not Rondeau. The percent of PAR penetrating the canopy was $log_{10}$ transformed.

| | Total stems (per $m^{-2}$) | | | Live stems (per $m^{-2}$) | | | PAR penetration (% incident light) | | |
|---|---|---|---|---|---|---|---|---|---|
| | df | F | P | df | F | P | df | F | P |
| Year | 1 | 1.258 | 0.265 | 1 | 0.883 | 0.350 | 1 | 2.476 | 0.120 |
| Location | 1 | 51.908 | < 0.001 | 1 | 0.023 | 0.880 | 1 | 7.085 | 0.009 |
| Year x Location | 1 | 4.286 | 0.0418 | 1 | 0.055 | 0.815 | 1 | 8.460 | 0.005 |
| Error | 78 | | | 78 | | | 78 | | |

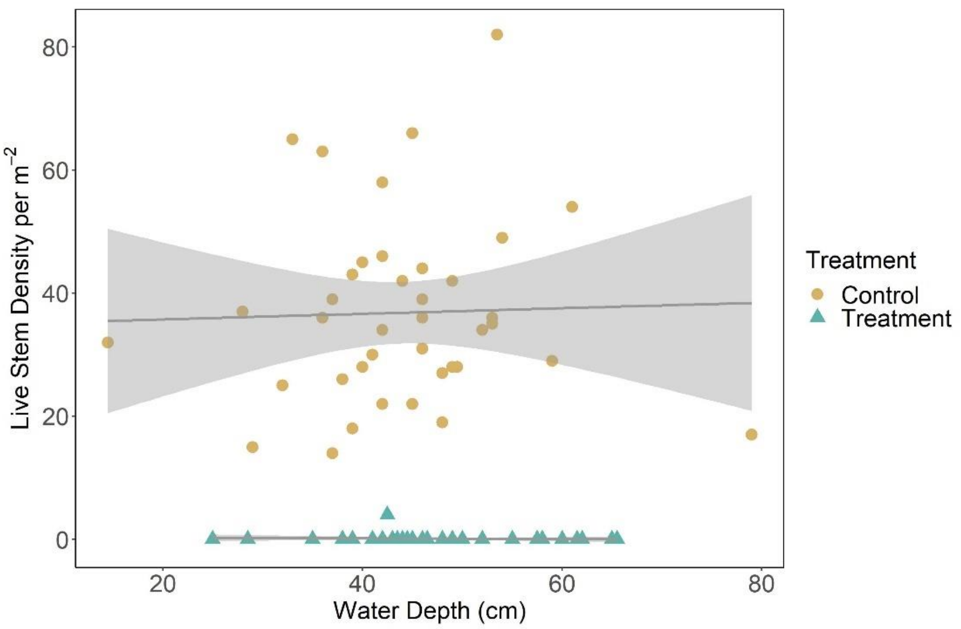


SM5. Glyphosate-based herbicide was effective at suppressing *P. australis* along the entire water depth gradient. The number of living *P. australis* stems (per $m^2$) in the control plots one year after treatment occurred were significantly higher than in the herbicide-treated plots and there was no difference in live *P. australis* stem density along the water depth gradient (cm). Shaded area represents 95% confidence intervals. Made with ggplot2 (Whickham 2016).

SM6. A significant difference in vegetation community composition arose between treatment and control plots after the herbicide was applied (i.e. the interaction term was significant). Thus, we conclude that treatment had an effect on the vegetation community structure. Results represent the average test statistics taken from 500 runs of a perMANOVA performed using stratified random sampling with replacement. Values in parentheses are the standard deviation.

| | df | Average Pseudo-F | Average p-value |
|---|---|---|---|
| Treatment | 1 | 170.20 (± 14.90) | 0.001 (< 0.001) |
| Year | 2 | 61.51 (± 6.13) | 0.001 (< 0.001) |
| Treatment x year | 2 | 44.77 (± 5.33) | 0.001 (< 0.001) |
| Residual | 222 | | |

SM7. The correlation coefficients (r) and coefficient of determination ($r^2$) of all the vectors in the 3D NMDS ordination for the control, treatment, and reference condition plots. Vectors with an $r^2 \geq 0.150$ are considered reasonably correlated with points and were included in figure 6.

| | Axis 1 | | Axis 2 | | Axis 3 | |
|---|---|---|---|---|---|---|
| Full name | r | $r^2$ | r | $r^2$ | r | $r^2$ |
| Water depth | -0.271 | 0.073 | 0.215 | 0.046 | 0.144 | 0.021 |
| Open water | -0.685 | 0.469 | -0.507 | 0.257 | -0.141 | 0.020 |
| Litter | -0.012 | 0.000 | 0.055 | 0.003 | -0.549 | 0.301 |
| Standing dead | -0.438 | 0.192 | -0.264 | 0.070 | 0.529 | 0.280 |
| *Calamagrostis canadensis* | -0.145 | 0.021 | -0.330 | 0.109 | -0.249 | 0.062 |
| *Campanula aparinoides* | -0.037 | 0.001 | -0.105 | 0.011 | -0.042 | 0.002 |
| *Calystegia sepium* | -0.070 | 0.005 | -0.090 | 0.008 | 0.055 | 0.003 |
| *Carex aquatilis* | -0.050 | 0.003 | -0.169 | 0.029 | -0.220 | 0.048 |
| *Carex buxbaumii* | -0.149 | 0.022 | -0.157 | 0.025 | -0.298 | 0.089 |
| *Carex crawei* | -0.102 | 0.010 | -0.128 | 0.016 | -0.058 | 0.003 |
| *Carex comosa* | 0.063 | 0.004 | 0.014 | 0.000 | -0.010 | 0.000 |
| *Carex lacustris* | -0.036 | 0.001 | 0.026 | 0.001 | 0.206 | 0.043 |
| *Carex lanuginosa* | -0.162 | 0.026 | -0.164 | 0.027 | -0.352 | 0.124 |
| *Carex lasiocarpa* | 0.017 | 0.000 | -0.119 | 0.014 | -0.134 | 0.018 |
| *Carex sartwellii* | -0.055 | 0.003 | -0.054 | 0.003 | -0.291 | 0.085 |
| *Carex* spp. | 0.055 | 0.003 | -0.004 | 0.000 | -0.045 | 0.002 |
| *Cladium mariscoides* | -0.130 | 0.017 | -0.109 | 0.012 | -0.392 | 0.154 |
| *Cornus stolonifera* | -0.146 | 0.021 | -0.172 | 0.030 | 0.005 | 0.000 |
| *Decodon verticillatus* | -0.010 | 0.000 | 0.058 | 0.003 | -0.009 | 0.000 |
| *Dulichium arundinaceum* | 0.091 | 0.008 | -0.006 | 0.000 | 0.007 | 0.000 |
| *Eleocharis palustris* | 0.173 | 0.030 | -0.023 | 0.001 | -0.017 | 0.000 |
| *Elodea canadensis* | 0.041 | 0.002 | -0.056 | 0.003 | -0.01 | 0.000 |
| *Eleocharis* spp. | -0.056 | 0.003 | -0.058 | 0.003 | -0.104 | 0.011 |
| *Equisetum fluviatile* | -0.071 | 0.005 | 0.183 | 0.034 | 0.017 | 0.000 |
| *Fontinalis* sp. | -0.103 | 0.011 | -0.028 | 0.001 | -0.016 | 0.000 |
| *Galium aparine* | 0.079 | 0.006 | -0.013 | 0.000 | -0.001 | 0.000 |
| *Hydrocharis morsus-ranae* | -0.384 | 0.147 | 0.866 | 0.749 | -0.056 | 0.003 |
| *Hypericum kalmianum* | -0.128 | 0.016 | -0.174 | 0.03 | -0.081 | 0.007 |
| *Iris versicolor* | -0.101 | 0.010 | -0.127 | 0.016 | -0.103 | 0.011 |
| *Juncus balticus* | -0.059 | 0.003 | -0.054 | 0.003 | -0.226 | 0.051 |
| *Lemna minor* | -0.042 | 0.002 | 0.137 | 0.019 | -0.073 | 0.005 |
| *Lysimachia thyrsiflora* | -0.145 | 0.021 | -0.156 | 0.024 | -0.156 | 0.024 |
| *Achillea millefolium* | -0.132 | 0.017 | -0.016 | 0.000 | -0.096 | 0.009 |
| *Myrioplyllum sibiricum* | -0.082 | 0.007 | 0.022 | 0.000 | -0.046 | 0.002 |
| *Myrioplyllum* spp. | -0.088 | 0.008 | 0.013 | 0.000 | 0.136 | 0.019 |
| *Nuphar variegatum* | 0.029 | 0.001 | 0.037 | 0.001 | -0.094 | 0.009 |
| *Nymphaea odorata* | 0.006 | 0.000 | 0.148 | 0.022 | -0.031 | 0.001 |
| *Phragmites australis* subsp. *australis* | 0.968 | 0.937 | -0.078 | 0.006 | 0.080 | 0.006 |

| | Axis 1 | | Axis 2 | | Axis 3 | |
|---|---|---|---|---|---|---|
| Full name | r | $r^2$ | r | $r^2$ | r | $r^2$ |
| *Polygonum amphibium* | 0.102 | 0.011 | -0.017 | 0.000 | 0.071 | 0.005 |
| *Polygonum* spp. | -0.133 | 0.018 | -0.071 | 0.005 | 0.198 | 0.039 |
| *Potamogeton* spp. | -0.083 | 0.007 | -0.067 | 0.004 | -0.058 | 0.003 |
| *Sagittaria* spp. | 0.054 | 0.003 | -0.028 | 0.001 | 0.026 | 0.001 |
| *Sagittaria latifolia* | -0.068 | 0.005 | 0.112 | 0.013 | -0.057 | 0.003 |
| *Scirpus acutus* | -0.028 | 0.001 | -0.079 | 0.006 | -0.007 | 0.000 |
| *Scirpus fluviatilis* | 0.074 | 0.006 | 0.004 | 0.000 | -0.047 | 0.002 |
| *Scirpus validus* | 0.046 | 0.002 | 0.01 | 0.000 | -0.004 | 0.000 |
| *Sparganium eurycarpum* | -0.072 | 0.005 | 0.209 | 0.044 | 0.026 | 0.001 |
| *Sparganium* spp. | -0.075 | 0.006 | 0.005 | 0.000 | -0.052 | 0.003 |
| *Spirodela polyrrhiza* | 0.065 | 0.004 | 0.01 | 0.000 | 0.005 | 0.000 |
| *Solonaceae* spp. | 0.046 | 0.002 | 0.03 | 0.001 | -0.027 | 0.001 |
| *Solidago* spp. | -0.113 | 0.013 | -0.143 | 0.02 | -0.051 | 0.003 |
| *Typha* spp. | -0.313 | 0.098 | 0.048 | 0.002 | 0.595 | 0.354 |
| *Typha angustifolia* | 0.220 | 0.048 | -0.016 | 0.000 | -0.125 | 0.016 |
| *Typha* x *glauca* | 0.180 | 0.032 | 0.065 | 0.004 | -0.231 | 0.053 |
| *Typha latifolia* | 0.060 | 0.004 | -0.022 | 0.001 | -0.049 | 0.002 |
| *Utricularia intermedia* | -0.111 | 0.012 | 0.004 | 0.000 | -0.222 | 0.049 |
| *Utricularia vulgaris* | -0.157 | 0.025 | -0.052 | 0.003 | -0.045 | 0.002 |
| *Unknown* | -0.093 | 0.009 | -0.069 | 0.005 | -0.083 | 0.007 |
| *Zizania palustris* | -0.085 | 0.007 | -0.115 | 0.013 | -0.033 | 0.001 |

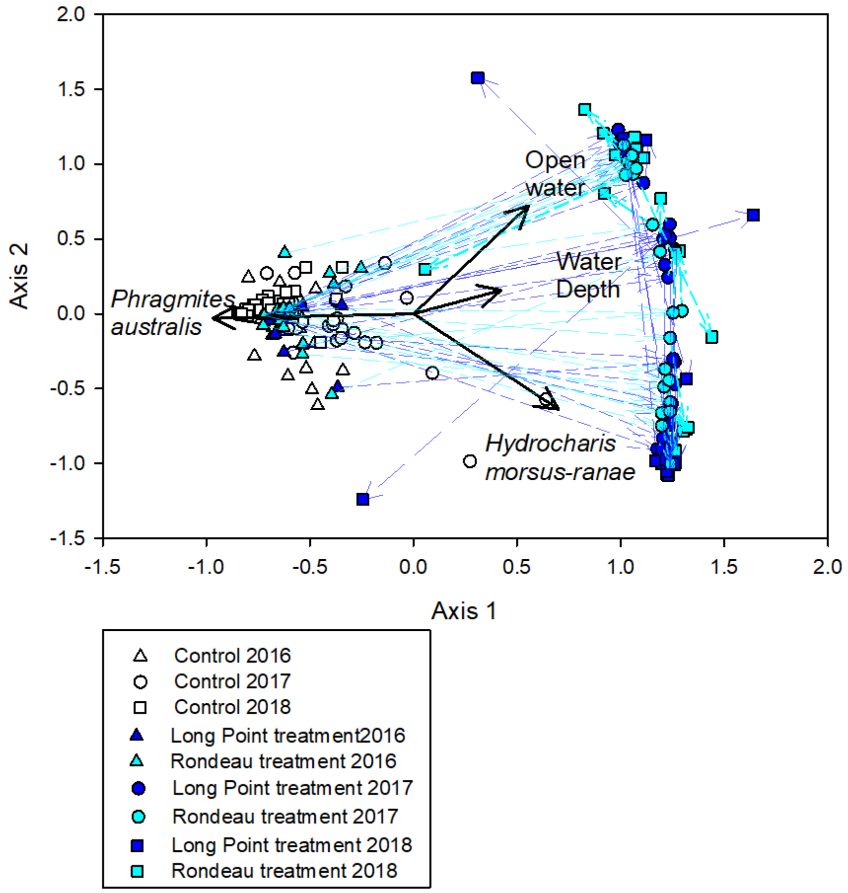


SM8. The final 2D NMDS ordination solution, using Bray-Curtis dissimilarity matrix, for the treatment and control plots before (2016) and two years after herbicide treatment. The ordination had a final stress of 10.348 with a final instability of < 0.001 after 63 iterations and explains 94.5% of the variance (Axis 1: 77.8%, Axis 2: 16.6%). Plots are symbolized by year and by whether they were located in Long Point, where secondary treatment occurred, or in Rondeau Provincial Park. Black vectors represent reasonably correlated ($r^2 \geq 0.150$) cover classes, and purple vectors represent the movement of plots between years.

SM9. The correlation coefficients (r) and coefficient of determination ($r^2$) of all the vectors in the 2D NMDS ordination for the control and treatment plots. Vectors with an $r^2 \geq 0.150$ are considered reasonably correlated with points.

| | Axis 1 | | Axis 2 | |
|---|---|---|---|---|
| Full name | r | $r^2$ | r | $r^2$ |
| Water depth | 0.423 | 0.179 | 0.158 | 0.025 |
| Open water | 0.555 | 0.308 | 0.722 | 0.521 |
| Litter | 0.072 | 0.005 | -0.012 | 0.000 |
| Standing dead | 0.366 | 0.134 | 0.302 | 0.091 |
| *Calmagrostis canadensis* | -0.145 | 0.021 | 0.014 | 0.000 |
| *Calystegia sepium* | -0.010 | 0.000 | 0.057 | 0.003 |
| *Carex aquatilis* | -0.138 | 0.019 | -0.012 | 0.000 |
| *Carex comosa* | -0.050 | 0.003 | -0.010 | 0.000 |
| *Carex lacustris* | -0.075 | 0.006 | 0.010 | 0.000 |
| *Carex lasiocarpa* | -0.076 | 0.006 | -0.016 | 0.000 |
| *Carex spp.* | -0.048 | 0.002 | -0.032 | 0.001 |
| *Decodon verticillatus* | 0.043 | 0.002 | 0.022 | 0.000 |
| *Dulichium arundinaceum* | -0.078 | 0.006 | 0.010 | 0.000 |
| *Eleocharis palustris* | -0.152 | 0.023 | -0.028 | 0.001 |
| *Eleocharis canadensis* | -0.035 | 0.001 | 0.063 | 0.004 |
| *Equisetum fluviatile* | 0.128 | 0.016 | -0.148 | 0.022 |
| *Fontinalis* spp. | 0.120 | 0.014 | 0.104 | 0.011 |
| *Galium aparine* | -0.070 | 0.005 | -0.004 | 0.000 |
| *Hydocharis morsus-ranae* | 0.700 | 0.491 | -0.637 | 0.406 |
| *Lemna minor* | 0.086 | 0.007 | -0.158 | 0.025 |
| *Lysimachia thrysiflora* | 0.085 | 0.007 | 0.012 | 0.000 |
| *Achillea millefolium* | 0.157 | 0.025 | 0.137 | 0.019 |
| *Myriophyllum sibiricum* | 0.103 | 0.011 | 0.050 | 0.003 |
| *Myriophyllum* spp. | 0.161 | 0.026 | 0.113 | 0.013 |
| *Nuphar variegatum* | -0.011 | 0.000 | -0.100 | 0.010 |
| *Nymphaea odorata* | 0.039 | 0.002 | -0.137 | 0.019 |
| *Phragmties australis* subsp. *australis* | -0.969 | 0.940 | -0.031 | 0.001 |
| *Polygonum amphibium* | -0.099 | 0.010 | 0.041 | 0.002 |
| *Potamogeton* spp. | 0.082 | 0.007 | 0.148 | 0.022 |
| *Sagittaria* spp. | -0.053 | 0.003 | 0.030 | 0.001 |
| *Sagittaria latifolia* | 0.130 | 0.017 | -0.041 | 0.002 |
| *Scirpus acutus* | 0.030 | 0.001 | 0.122 | 0.015 |
| *Scirpus fluviatilis* | -0.066 | 0.004 | 0.029 | 0.001 |
| *Scirpus validus* | -0.038 | 0.001 | -0.008 | 0.000 |
| *Sparganium eurycarpum* | 0.134 | 0.018 | -0.179 | 0.032 |
| *Sparganium* spp. | 0.092 | 0.008 | 0.065 | 0.004 |
| *Spirodela polyrrhiza* | -0.056 | 0.003 | -0.010 | 0.000 |
| *Solonaceae* spp. | -0.032 | 0.001 | -0.031 | 0.001 |
| *Typha* spp. | 0.291 | 0.085 | -0.058 | 0.003 |
| *Typha angustifolia* | -0.185 | 0.034 | -0.043 | 0.002 |
| *Typha* x *glauca* | -0.122 | 0.015 | -0.175 | 0.031 |
| *Typha latifolia* | -0.048 | 0.002 | 0.028 | 0.001 |
| *Utricularia intermedia* | 0.125 | 0.016 | -0.049 | 0.002 |
| *Utricularia vulgaris* | 0.175 | 0.031 | 0.214 | 0.046 |

| Full name | Axis 1 | | Axis 2 | |
|---|---|---|---|---|
| | r | $r^2$ | r | $r^2$ |
| Unknown species | 0.068 | 0.005 | 0.026 | 0.001 |
| *Zizania palustris* | 0.081 | 0.007 | 0.209 | 0.044 |

SM10. Movement of treatment plots as a proportion of the total Euclidean distance between a reference condition centroid and the control centroid. Reference conditions for each plot were selected by choosing the reference condition (emergent marsh or meadow marsh) it was closest to in 2018. The proportion of that distance was then calculated as $((Dist_{Ref})/(Dist_{Ref} + Dist_{Con})*100)$, and as such a smaller value in 2018 than in 2017 would represent a plot moving towards its reference condition. This was true for only 16 of the treatment plots, indicating that most are not moving towards a composition that reflects the reference vegetation communities.

| Plot | Park | Reference | 2017 | 2018 | If 2018 < 2017 |
|---|---|---|---|---|---|
| LPT05 | Long Point | Emergent | 45.6 | 50.0 | FALSE |
| LPT06 | Long Point | Emergent | 51.1 | 50.3 | TRUE |
| LPT07 | Long Point | Emergent | 48.6 | 45.8 | TRUE |
| LPT08 | Long Point | Emergent | 46.8 | 43.9 | TRUE |
| LPT09 | Long Point | Meadow | 18.5 | 27.4 | FALSE |
| LPT10 | Long Point | Emergent | 41.9 | 24.0 | TRUE |
| LPT11 | Long Point | Emergent | 41.5 | 50.3 | FALSE |
| LPT12 | Long Point | Emergent | 42.8 | 50.0 | FALSE |
| LPT13 | Long Point | Emergent | 44.6 | 49.6 | FALSE |
| LPT14 | Long Point | Emergent | 48.5 | 49.1 | FALSE |
| LPT15 | Long Point | Emergent | 42.3 | 46.3 | FALSE |
| LPT16 | Long Point | Emergent | 41.7 | 46.7 | FALSE |
| LPT17 | Long Point | Emergent | 42.4 | 53.1 | FALSE |
| LPT18 | Long Point | Emergent | 41.6 | 49.2 | FALSE |
| LPT19 | Long Point | Meadow | 36.1 | 50.5 | FALSE |
| LPT20 | Long Point | Emergent | 35.7 | 25.6 | TRUE |
| LPT41 | Long Point | Emergent | 40.8 | 40.1 | TRUE |
| LPT42 | Long Point | Emergent | 46.4 | 47.3 | FALSE |
| LPT43 | Long Point | Emergent | 49.1 | 49.7 | FALSE |
| LPT44 | Long Point | Emergent | 40.2 | 45.9 | FALSE |
| RPC16 | Long Point | Meadow | 29.9 | 29.8 | TRUE |
| RPT21 | Long Point | Emergent | 47.8 | 49.3 | FALSE |
| RPT22 | Long Point | Emergent | 46.2 | 48.6 | FALSE |
| RPT23 | Long Point | Emergent | 47.4 | 48.6 | FALSE |
| RPT24 | Long Point | Emergent | 44.8 | 48.6 | FALSE |
| RPT25 | Long Point | Meadow | 24.4 | 22.3 | TRUE |
| RPT26 | Long Point | Emergent | 37.5 | 25.8 | TRUE |
| RPT27 | Long Point | Meadow | 12.3 | 15.5 | FALSE |
| RPT28 | Long Point | Emergent | 44.4 | 43.1 | TRUE |
| RPT29 | Long Point | Emergent | 46.4 | 44.3 | TRUE |
| RPT30 | Long Point | Emergent | 45.2 | 44.3 | TRUE |
| RPT31 | Long Point | Emergent | 44.5 | 48.6 | FALSE |
| RPT32 | Long Point | Meadow | 39.8 | 33.8 | TRUE |
| RPT33 | Long Point | Meadow | 18.8 | 22.2 | FALSE |
| RPT34 | Long Point | Meadow | 19.6 | 43.5 | FALSE |
| RPT35 | Long Point | Meadow | 25.2 | 24.7 | TRUE |

| Plot | Park | Reference | 2017 | 2018 | If 2018 < 2017 |
|---|---|---|---|---|---|
| RPT36 | Long Point | Meadow | 23.2 | 25.2 | FALSE |
| RPT37 | Long Point | Meadow | 23.9 | 28.1 | FALSE |
| RPT38 | Long Point | Meadow | 23.9 | 21.4 | TRUE |
| RPT39 | Long Point | Meadow | 21.8 | 57.0 | FALSE |
| RPT40 | Long Point | Meadow | 39.3 | 32.8 | TRUE |
